# A transcription factor Abf1 facilitates ORC binding onto the *Saccharomyces cerevisiae* replication origin via histone acetylase Gcn5

**DOI:** 10.1101/583310

**Authors:** Hidetsugu Kohzaki, Yota Murakami

**Author notes:** Faculty of Nursing, Shumei University, Daigaku-machi 1-1, Yachiyo, Chiba, Japan 276-0003 (Hidetsugu Kohzaki). Department of Chemistry, Faculty of Science, Hokkaido University, Kita-ku, Sapporo, Hokkaido, Japan 060-0810 (Yota Murakami). Corresponding author: Hidetsugu Kohzaki, Department of Viral Oncology, Institute for Virus Research, Kyoto University, shogoin, Sakyo-ku, Kyoto 606-8507, Japan. We dedicate this work to Dr. Yasuo Kawasaki and Dr. Yuko Yamaguchi-Iwai.

## Abstract

Chromatin structure has been implicated in the regulation of DNA replication but the molecular mechanism involved is unclear. In this study, we observed that binding of the transcription factor Abf1 to the replication origin *ARS1* facilitated the association of the origin recognition complex (ORC) with *ARS1* using genetic interaction analysis and ChIP assay. The histone acetyltransferases (HATs), Gcn5 and Esa1, were also loaded onto *ARS1* in an Abf1 site-dependent manner, where they were then responsible for acetylating histone H3 lysine 18 (H3K18) and histone H4 lysine 12 (H4K12), respectively. Interestingly, Abf1 interacted with Gcn5, while ORC interacted with Esa1. Indeed the B3 element showed genetic interaction with Gcn5 and Rpd3 not with Esa1, Act3 and Tra1.

These data suggest that Gcn5, which is recruited by Abf1, alters chromatin structure via histone acetylation and facilitates the loading of ORC. We therefore propose that transcription factors regulate chromatin structure at replication origins by recruiting chromatin-modifying proteins, such as HATs, to load the initiator.

## Introduction

Initiation of DNA replication is highly regulated for precise genome duplication and maintenance of genome integirity. Replicon model proposed that DNA replication starts when a DNA binding protein or protein complex called an initiator binds to a defined chromosome region replicator (Jacob et al, 1964). This model seems to be applicable to almost all organisms so far analyzed. DNA replication in eukaryote cells initiates from large numbers of replication origins distributed on multiple chromosome.

In *Saccharomyces cerevisiae*, specific ARS (autonomously replicating sequence) element was identified on the basis of their ability to direct the autonomous replication of cloned plasmid DNA (Newlon and Theis, 1993; Shirahige, et al. 1993; Huberman 1999). ARS contains a consensus sequence (ARS consensus sequence or ACS or A element) (Newlon and Theis, 1993). The eukaryotic initiator protein, origin recognition complex (Orc) is first isolated from *Saccharomyces cerevisiae* as a six subunit ACS binding protein complex (Bell et al., 1993) and shown to be essential for the initiation of DNA replication. However, an ACS alone is not sufficient for origin function and this sequence is much more abundant than the number of the ORC binding sites or the functional replication origins on the genome (Marahrens and Stillman, 1992; River and Rine, 1992; Newlon and Theis, 1993; Rao et al., 1994). In addition to an ACS, many ARSs contain at least one A/T–rich region of DNA that is thought to act as a DNA unwinding element (B2 element). Though this element is important for ARS activity, it is not always required for ORC binding.

The chromosome replication cycle proceeds through multiple steps (Masumoto et al., 2000; Bell and Dutta, 2002; Bielinsky and Gerbi, 2001; Kearsey and Cotterill, 2003; Méndez and Stillman, 2003; Diffley, 2004; Cevetic and Walter, 2005; Kohzaki and Murakami, 2005; Moyer et al, 2006). The Orc is shown to stably associate with ARS during cell cycle progression. The putative replicative helicase complex, Mcm2-7 (Ishimi 1997; Shechter et al, 2004; Takahashi et al, 2005), is recruited onto Orc-bound origins with asist of Cdc6 and Cdt1 (Whittaker et al, 2000; Tanaka and Diffley, 2002) resulting in formation of pre-replicative complex (pre-RC). The pre-RC is activated by Cdc7-Dbf4 kinase (DDK) and Cdks. DDK phosphorylates the subunits of Mcm2-7 (Jares et al., 2000; Sclafani, 2000; Masai, et al., 2006; Sheu and Stillman, 2006) and Cdc45 (Jare and Blow, 2000; Zou and Stillman, 2000) leaded to changing the conformation of the complex to facilitate the loading of subsequent factors. The CDK phosphorylates Sld2 and Sld3 to form CMG (Cdc45-Mcm2-7-GINS) complex which leads to assemble DNA replication fork complex containing PCNA, RF-C, PR-A and replicative DNA polymerases, α, δ and ε (Moyer et al, 2006).

Large body of evidence shows the correlation between transcription and DNA replication (Hyrien et al., 1995; Sasaki, 1999; Gilbert, 2001; Méchali, 2001; Lunyak et al., 2002; Schübeler et al., 2002; MacAlpine et al., 2004; Danis et al 2004; Kohzaki and Murakami, 2005). In fact, the transcription factors regulate the chromosomal DNA replication in DNA virus, yeast and mammals. In SV40 and Polyomavirus replication, transcription factors bound to the vicinity of origin stimulate initiation of DNA replication through multiple mechanisms. GAL4-VP16, Gal4, p53 and BPV E2 stimulated DNA replication through interaction with single stranded DNA binding protein, RP-A (Kohzaki and Murakami, 2005). Runx1 stimulates DNA replication by tether origin to nuclear matrix where viral DNA replication takes place (Chen et al., 1998; Murakami et al., 2007). c-Jun recruited viral initiator, Large T antigen (Tg) to replication origin for the formation of origin-Tg initiation complex (Ito et al., 1996).

In ARS1 of Saccharomyces cerevisiae, the acidic activation domains of Gal4, p53, VP16, c-Jun, BRCA1 and replication activation domain of Runx1) have been shown to regulate replication activity when they were tethered to ARS1 (Marahrens and Stillman, 1992; Li et al., 1998; Hu et al., 1999; Kohzaki et al, 1999). In our previous study (Kohzaki et al. 1999), we showed that transcription factors regulate the activity of the *ARS* in a context-dependent manner, suggesting that the chromatin structure around replication origins may play an important role in replication, as has been suggested in organisms other than *S. cerevisiae* (Gerbi and Bielinsky 2002). In addition, it has been shown that the chromatin structure, particularly the state of histone acetylation, surrounding replication origins, can regulate the timing of origin firing (Vogelauer et al. 2002). However, the molecular mechanisms by which transcription factors or chromatin structure regulate cellular DNA replication are not yet understood (Raghuraman et al. 2001; Kim et al. 2003; Wyrick et al. 2003).

In this report, we showed that the B3 element of *ARS1* that is a binding sites for transcription factor Abf1 facilitates Orc binding onto origin using ChIP assay and genetic analyses. Level of acetylation of histone H3 lysine 18 (H3K18) and histone H4 lysin 12 (H4K2), which are mark for active chromatin, was significantly decreased without B3 element. Transcription factors, LexA-VP16 and Gcn4 can rescue Orc loading as well as histon actylation. Chromatin inmmunoprecipitation and genetic interaction experiments showed that Gcn5 that acetylaes H3K18 associate with ARS1 through Abf1 bound to B3, while Esa1 that acetylate H4K12 located onto *ARS1* through Orc. These data suggested that Abf1 recruits Gcn5 to provide an active chromatin structure for ORC binding and then Orc recruits Esa1 for subsequent replication steps.

## Results

To clarify the mechanism by which transcription factors regulate the activity of replication origins, we first analyzed the effect of various replication-related mutants (see Supplemental information) on to the replication of ARS1 plasmids harboring mutations in B3 or B2 element (Table S4). We prepared three B3 mutants, B3/mB3, B3/Gal4 and B3/LexA as shown in Fig.S2. In the B3 mutant, B3/mB3, the B3 element was replaced with the point mutation in ABF1 binding site (Marahrens and Stillman 1992; Kohzaki et al. 1999). In the B3 mutants, B3/Gal4 and B3/LexA, the B3 element was replaced with a Gal4 binding site and a LexA binding site, respectively (Marahrens and Stillman 1992; Kohzaki et al. 1999). We also prepared the mB2 mutant, in which B2 element is changed for Xho linker in Fig.S2. Since B3 is a binding site for transcription factor, Abf1 and B2 is corresponding to DNA unwinding element, both of which are required for efficient ARS1 plasmid replication, mutations in B2 or B3 caused instability of the ARS plasmid (Marahrens and Stillman 1992). Mutations of genes coding MCM proteins (mcm5, mcm7), DNA polymerases (pol1, cdc17), CDC28 (cdc28as-1), GINS (dpb2, dpb11), single stranded DNA binding protein (rfa2), RFC (cdc44) and polymerase-loading protein (cdc45, JET1) increased the instability of the B2-mutant ARS1 plasmid but did not affect that of the B3 mutant plasmid. In contrast, mutations in genes coding ORC (orc1, orc2) or ABF1 (abf1) specifically increased the instability of the B3 mutant plasmid. Interestingly, mutation in of a gene coding CDC6 (cdc6-1) did not affect the instability of the B2 and B3-mutant ARS1 plasmid (Table. S4). These data showed that B3 element genetically interacts with ORC and ABF1. The genetic interaction between B2 element and various replication mutants including DNA polymerases and MCM is consistent with the notion that B2 element is the site where DNA is unwound and replication fork complex is formed.

Next we investigated which step in the initiation of replication is regulated by the transcription factors binding to *ARS1* by comparing the ability of wild-type and mutant *ARS1* to bind to the replication proteins using the ChIP assay. We used yeast strains bearing mutations in the B2 (mB2 in Fig.S2) or the B3 (B3/mB3, B3/Gal4 and B3/LexA in Fig.S2) elements of chromosomal *ARS1* (Marahrens and Stillman 1992) (Fig. S2).

We first analyzed the association of ORC with the chromosomal origins *ARS1* and *ARS305*, both of which are strong ARSs (Fig. S3) by ChIP assay for tagged Orc subunits (myc-tagged ORC2 or HA-tagged ORC1) with cells arrested at the late G1, early S, G1/S or M phases. In all phases tested, the HA-tagged Orc1 and the myc-tagged Orc2 subunit associated with both origin sequences, but not with the non-origin sequence, *CYC1(*lane 5 in each panel and S3A-E). The B3 mutation in *ARS1* reduced the association of ORC with *ARS1* (Fig. S3F), but not with *ARS305*, in the late G1, early S and M phases (lane 7 in each panel and S3G). As previously reported (Bell and Dutta 2002; Kearsey and Cotterill 2003; Méndez and Stillman 2003; Diffley 2004; Moyer et al. 2006), the Mcm complex, Cdc45, Rfa, Rfc, Pol2 and DNA polymeraseα-primase are sequentially loaded onto the origins, as shown by the timing of appearance of ChIP signals of HA-tagged Mcm4, a subunit of the Mcm complex (Fig. 1A, lane 4); HA-tagged Cdc45 (Fig. S3I), myc-tagged Rfa1, a subunit of Rfa (Fig. 1B, lane 5) HA-tagged Pol2 (Fig. S3H) and myc-tagged Pri1, a subunit of the DNA polymerase α-primase (pol α-primase) complex (Fig. 1C, lane 8). Loading of these proteins onto the mutant *ARS1* (B3/LexA) was not detected (Fig. S3, lane 7; Fig. 1A, lane 6; Fig. 1B, lane 7; Fig. 1C, lane 10 in each panel, Fig S3), while loading onto *ARS305* was unaffected in the same strain. Since the loading of thess replication factors depend on the loading of ORC (Bell and Dutta 2002; Kearsey and Cotterill 2003; Méndez and Stillman 2003; Diffley 2004; Moyer et al. 2006), these results show that the B3 element is of primary importance in ORC-loading onto *ARS1* and that mutation of this element disrupts subsequent loading or replication factors. Exogenous expression of LexA-VP16 fusion protein could support ORC-loading onto ARS1 (B3/LexA), but expression of LexA DNA binding domain alone could not (Supplemental Discussion and Fig. S4). These results indicated that the primary function of transcription factors during replication is the loading of ORC onto the replication origins, which is consistent with the genetic interaction between B3 element and ORC (Figure S4).

**Figure 1.**
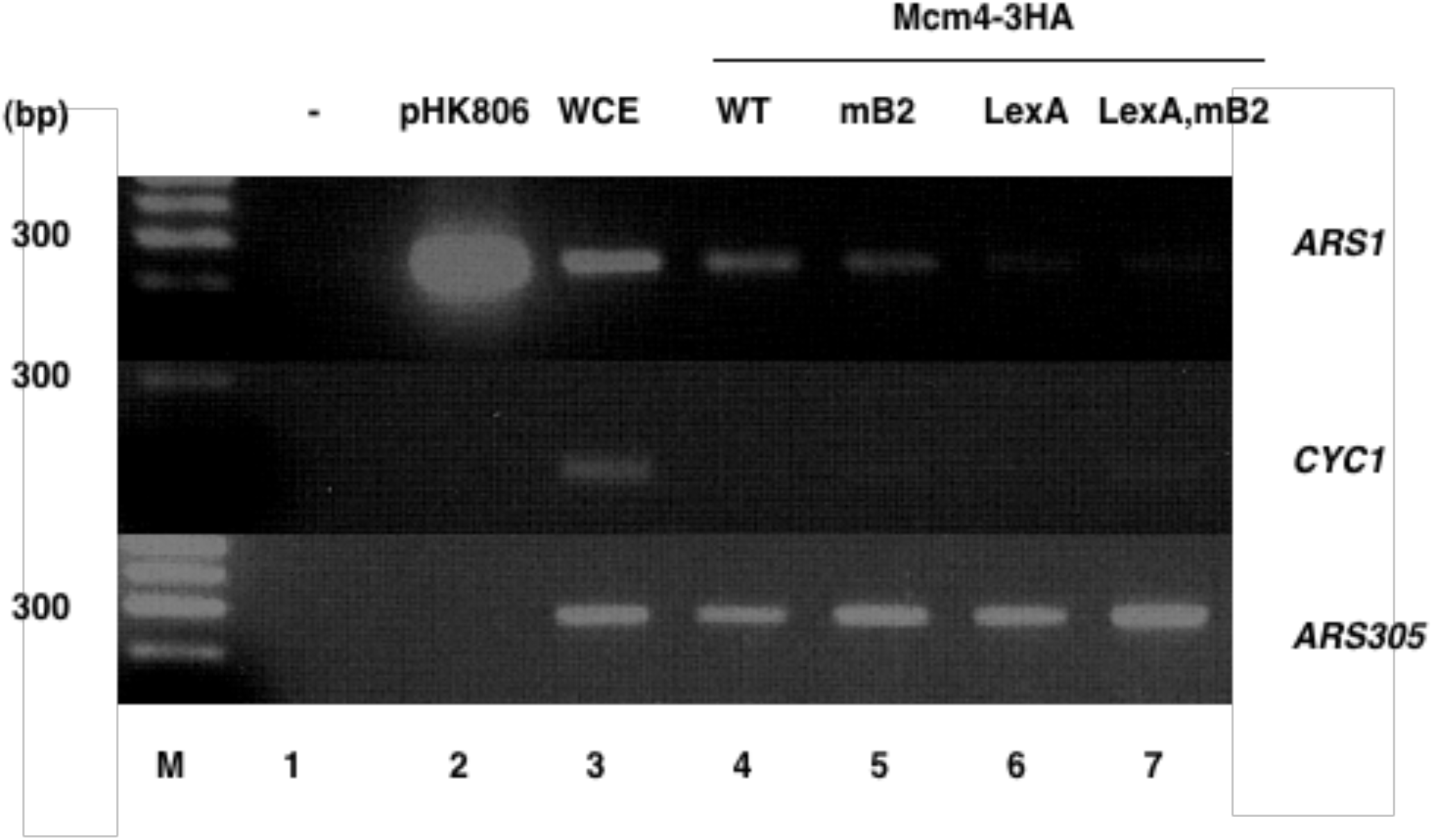
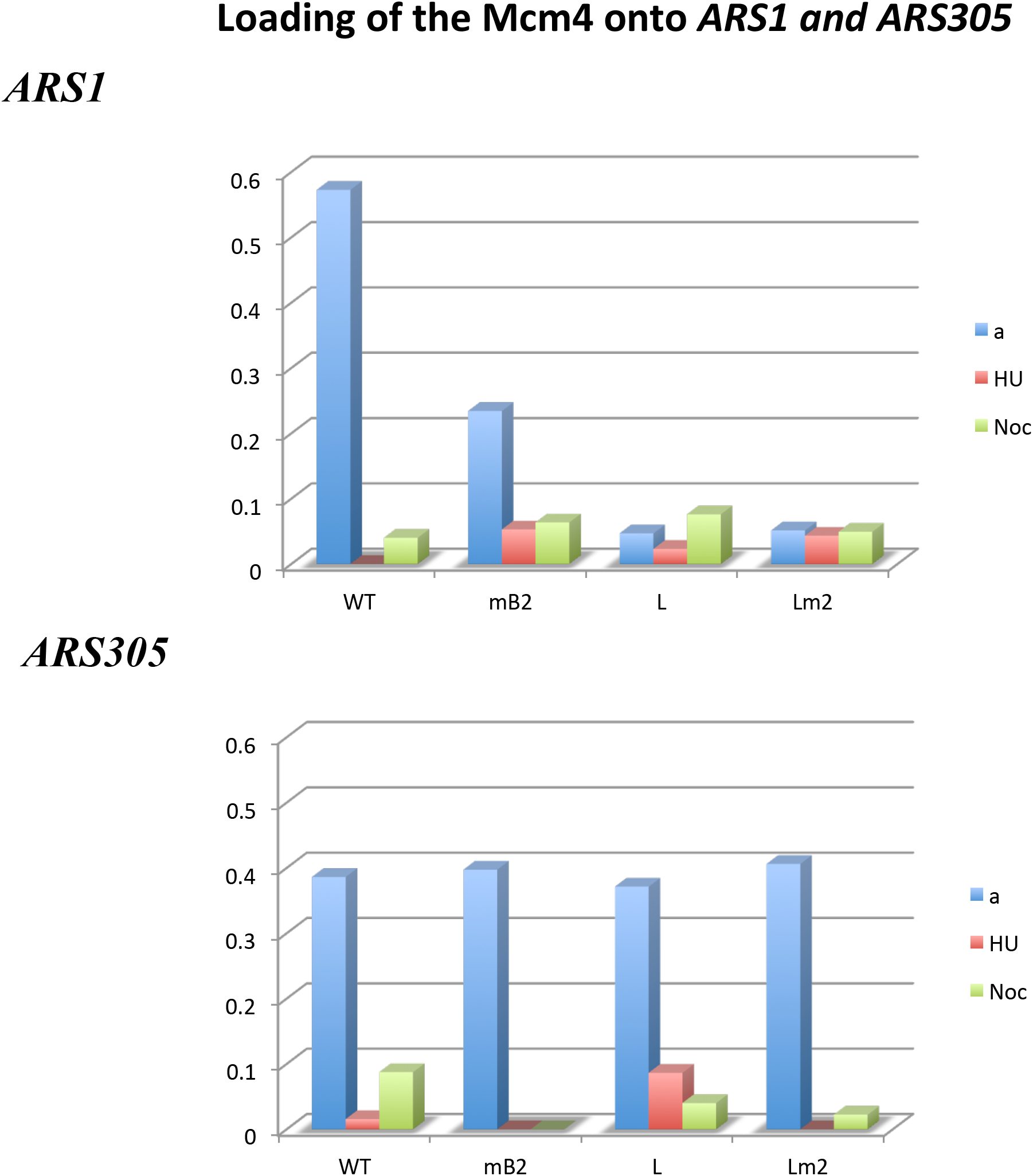
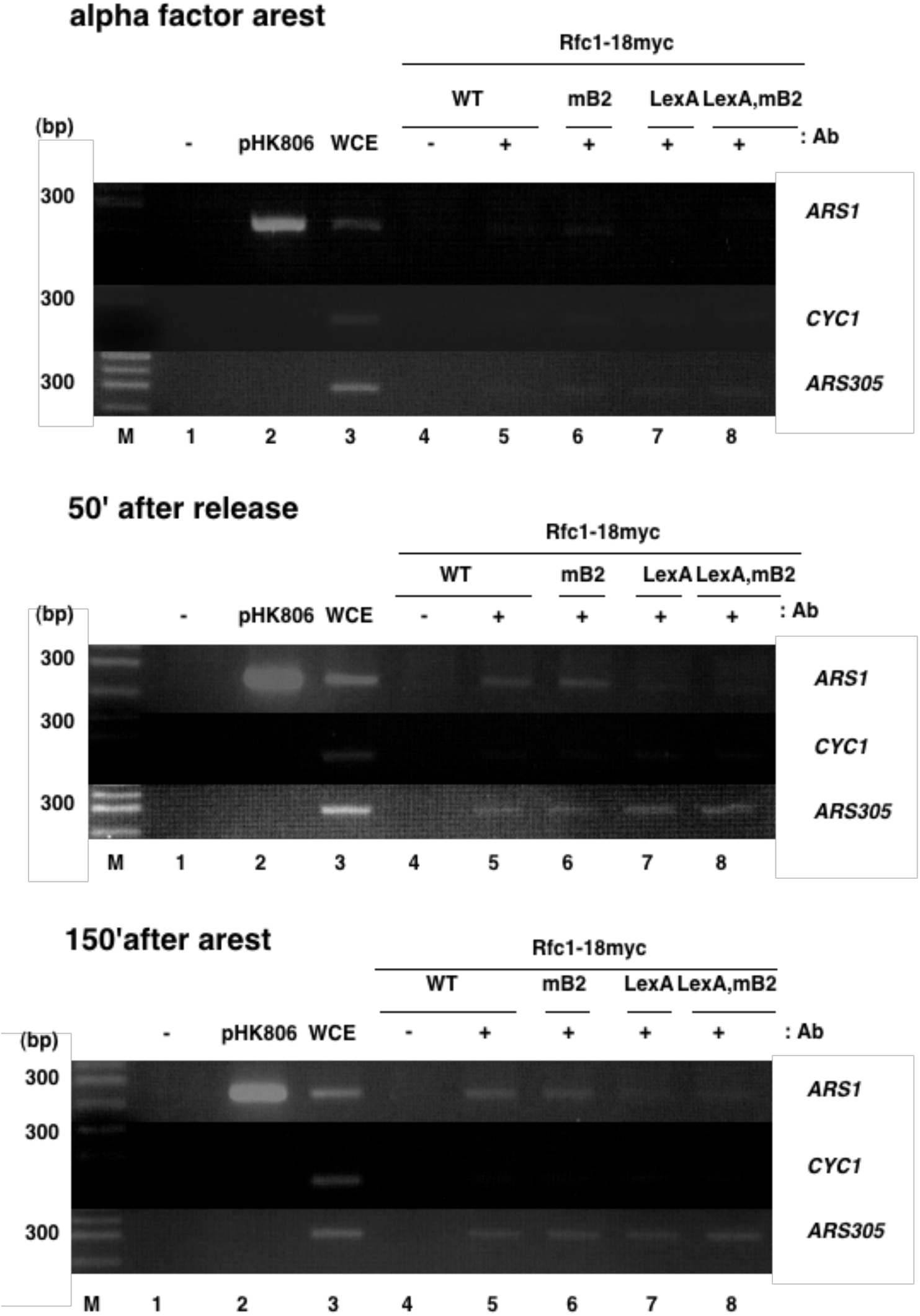
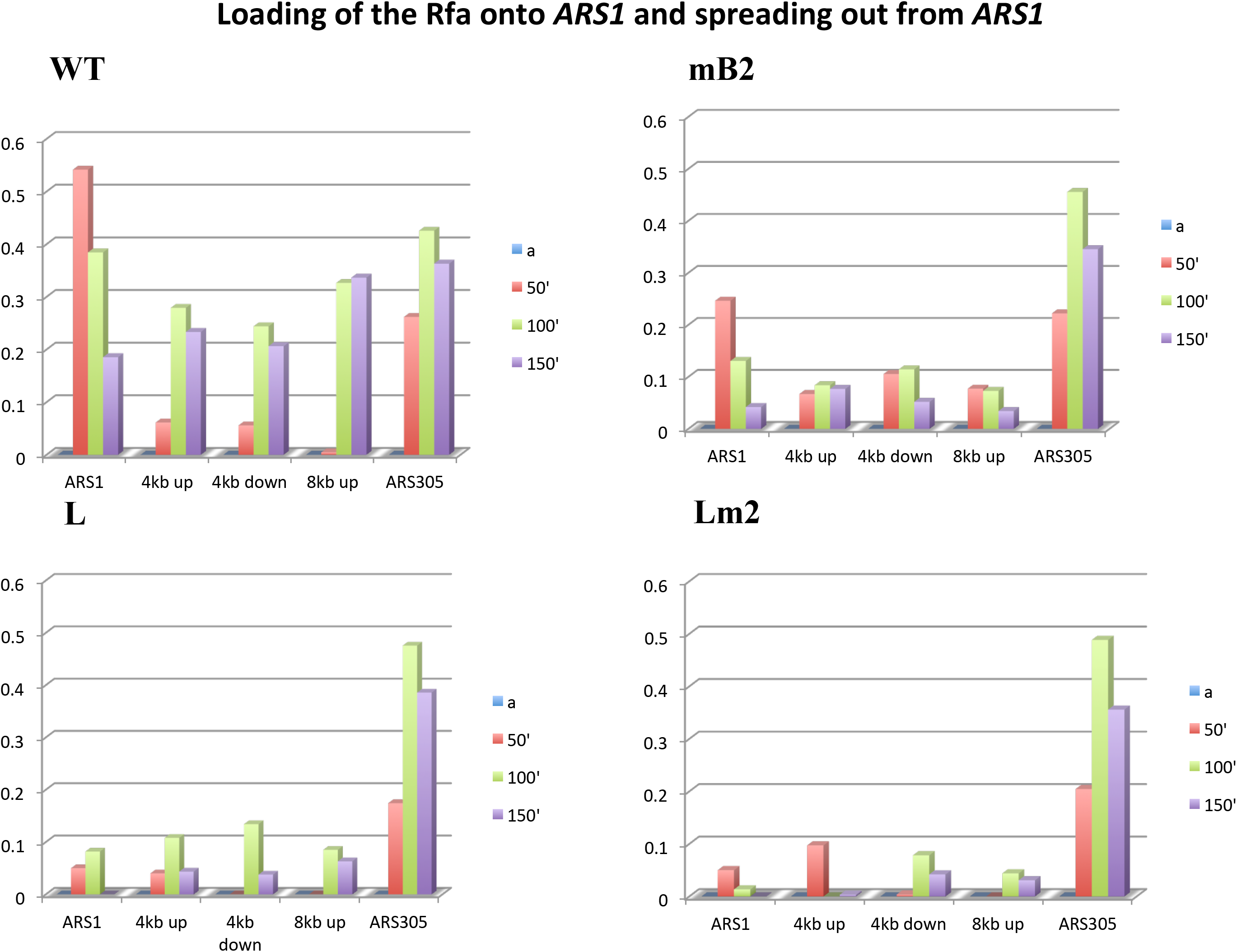
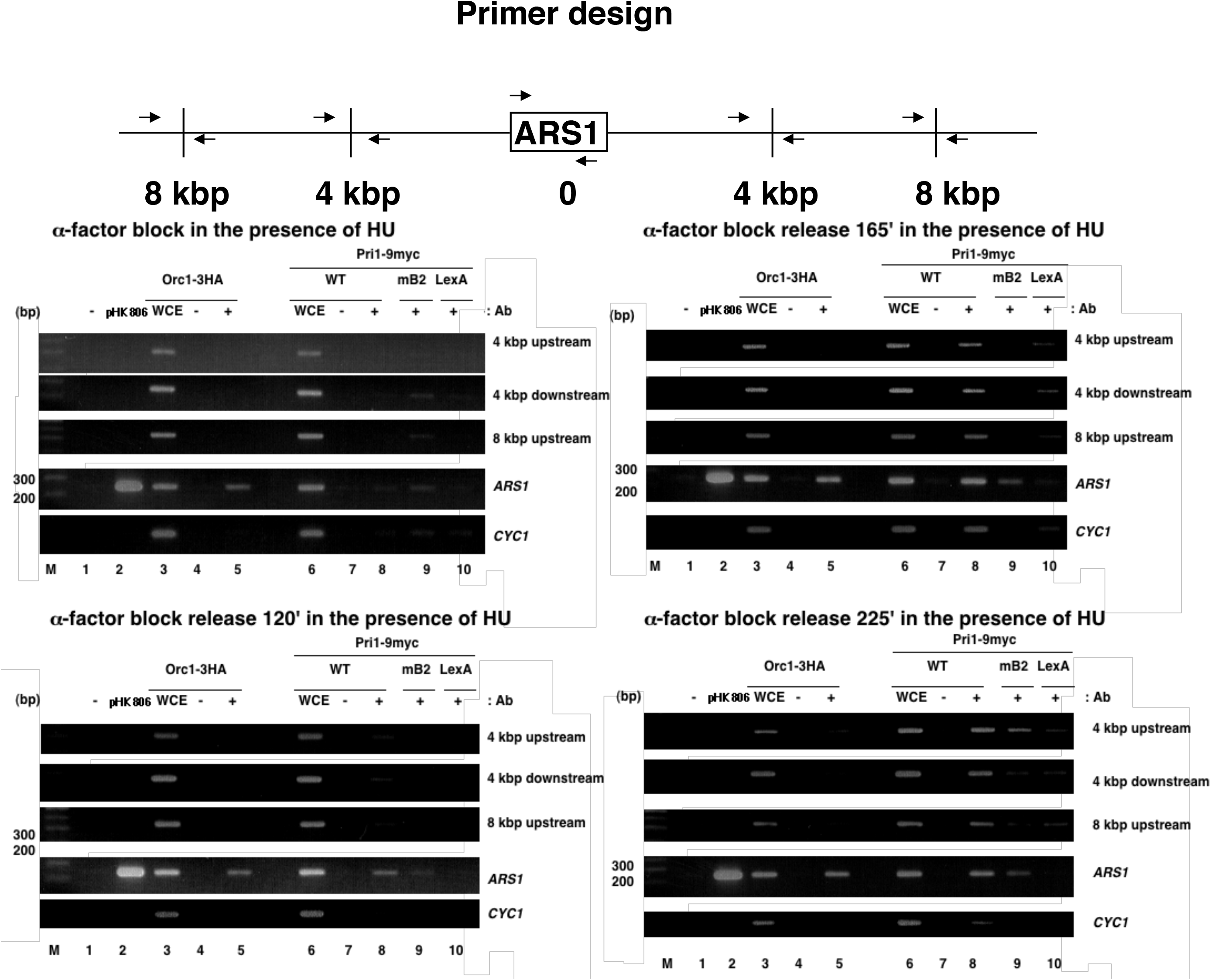
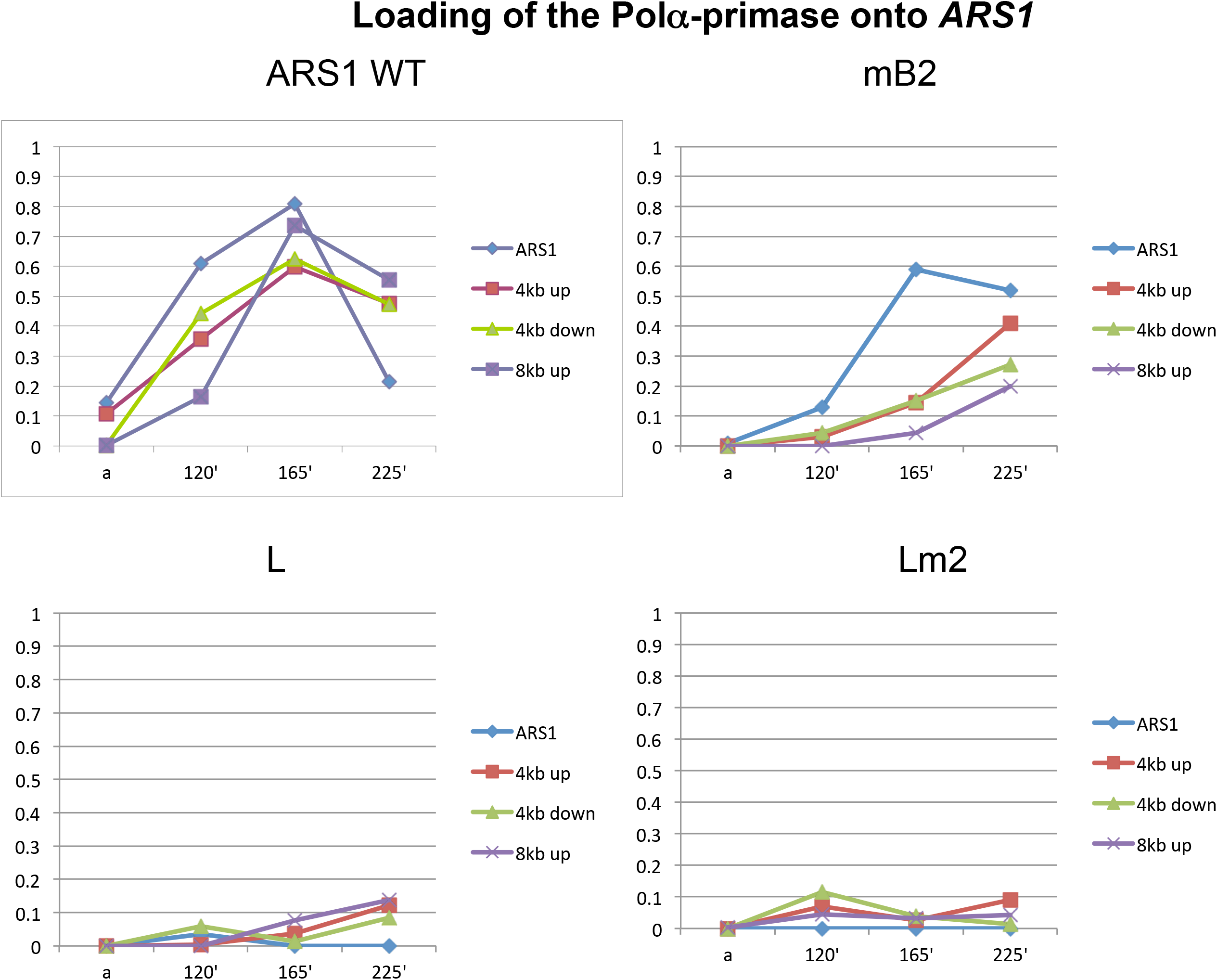
Abf1 regulates ORC loading onto *ARS1*. (*A*) Loading of the MCM complex onto *ARS1*. Cells harboring HA-tagged Mcm4 (Mcm4-3HA), which is a subunit of the Mcm complex, and the *ARS1* mutations indicated in the figure were synchronized with α-factor in late G1. ChIP assays were performed with anti-HA antibodies as described in the Methods section. The experiments were repeated three times, with similar results. Yeast strains used were YHM08, YHK007, YHK008 and YHK009 (Table S2). M, molecular weight marker. (*B*) The amounts of ARS1 WT PCR products obtained with ChIP assay relative to that obtained from mB2, L and Lm2 (using T testing, P<0.05) but not in ARS305 in A. (*C*) Cells harboring myc-tagged Rfc1 (Rfc1-18myc) and the *ARS1* mutations indicated in the figure, were synchronized at late G1 with α-factor and released from the block at 37°C in the presence of HU in order to block the cells in the next G1 phase. Samples were collected for the ChIP assay at the indicated time points. ChIP assays were performed with anti-HA antibodies as described in the Methods section. The experiments were repeated three times, with similar results. Yeast strains used were YHM014, YHK004, YHK005 and YHK006 (Table S2). (*D*) The amounts of ARS1 WT PCR products obtained with ChIP assay relative to that obtained from mB2, B3/LexA and B3/LexA, mB2(Lm2) (using T testing, P<0.05) but not in ARS305. (*E*) Loading of pol α-primase onto *ARS1*. (upper panel) Locations of PCR primers on the chromosome (arrows). (lower panel) Cells harboring myc-tagged Pri1 (Pri1-9myc), which is a subunit of pol α-primase, and the mutations indicated in *ARS1*, were synchronized in late G1 with α-factor and released from the block at 37°C in the presence of HU in order to block the cells in the next G1 phase. Samples were collected for ChIP assays at the indicated time points. ChIP assays were performed with anti-myc antibodies as described in the Methods section. PCR analysis using primers designed to amplify the non-origin sequence *CYC1* (upper panel) was performed on the precipitated DNA and the products were analyzed by agarose gel electrophoresis and EtBr staining. The experiments were repeated twice, with similar results. Yeast strains used were YHM07, YHK013, YHK001 and YHK002 (Table S2). M, molecular weight marker. (*F*) The amounts of ARS1 WT PCR products obtained with ChIP assay relative to that obtained from mB2, B3/L and B3/LexA,mB2(Lm2) (using T testing, P<0.05) but not in ARS305.

In contrast, we found that the loading and the movment of Pri1 was delayed in the presence of the B2 mutation, although the loading of other factors (Orc2) was not affected. We also found that the loading of Cdc45, Rfa, Mcm and Pol2 delayed in the presence of the B2 mutation (Figure 1A,B and S3H, I). This is consistent with these factors genetically interact with B2 element (Figure S4). These results suggest that the B2 element is involved in the formation of replication forks (see Fig. 1, Fig. S3H, I and the Supplemental Discussion).

One of the functions of transcription factors during transcriptional activation is changing the chromatin structure around promoters (Workman 2006; Shahbazian and Grunstein 2007) and the relationship between histone acetylation and active chromatin has been well documented (Workman 2006; Shahbazian and Grunstein 2007). Therefore, we examined the effect of the B3 mutation on histone acetylation around *ARS1*. The ChIP assay was performed using the anti-H4K12Ac and anti-H3K18Ac antibodies (Suka et al. 2001; Vogelauer et al. 2002) and showed that both the chromosomal and plasmid *ARS1* and *ARS305* sequences were acetylated at histone H4K12 and H4K18 in cells either growing asynchronously or blocked at late G1 by α-factor (Fig. 2A, lanes 5 and 11; Fig. S5, lane 5; Fig. 2B, lanes 5 and 11; Fig. 2C; Fig. S6, lane 5). Mutation of the B3 element, but not of B2, decreased the acetylation of both histones on *ARS1* to undetectable levels but did not affect acetylation on *ARS305* (Fig. 2A, lanes 7 and 13; Fig. S5, lane 6; Fig.2B, lanes 7 and 13; Fig. S6, lane 6). These results showed that active *ARS* sequences are hyperacetylated, a state that represents “active chromatin” and that, in *ARS1*, the B3 mutation changed the acetylation state to the hypoacetylated state, which represents “inactive” chromatin.

**Figure 2.**
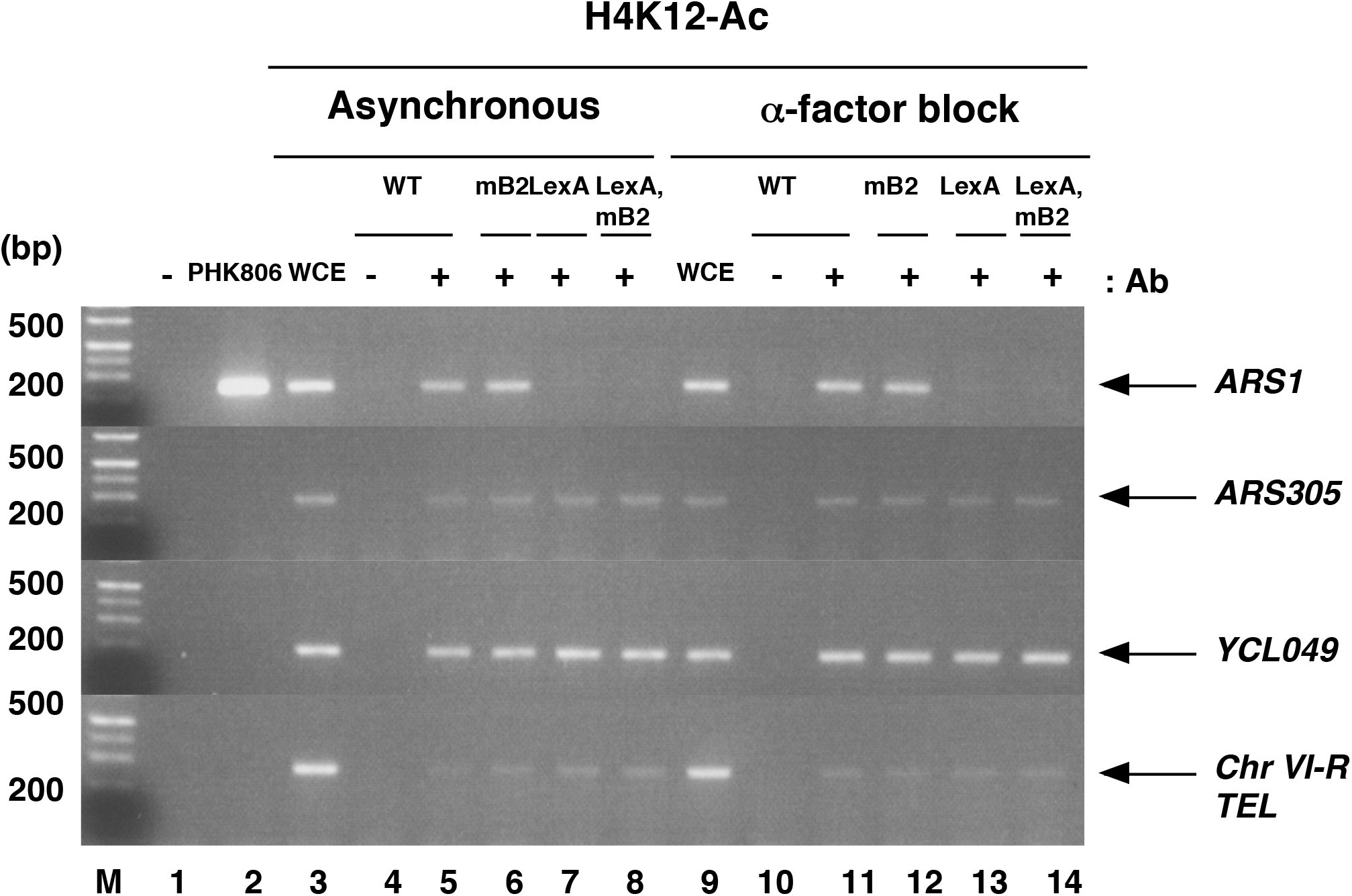
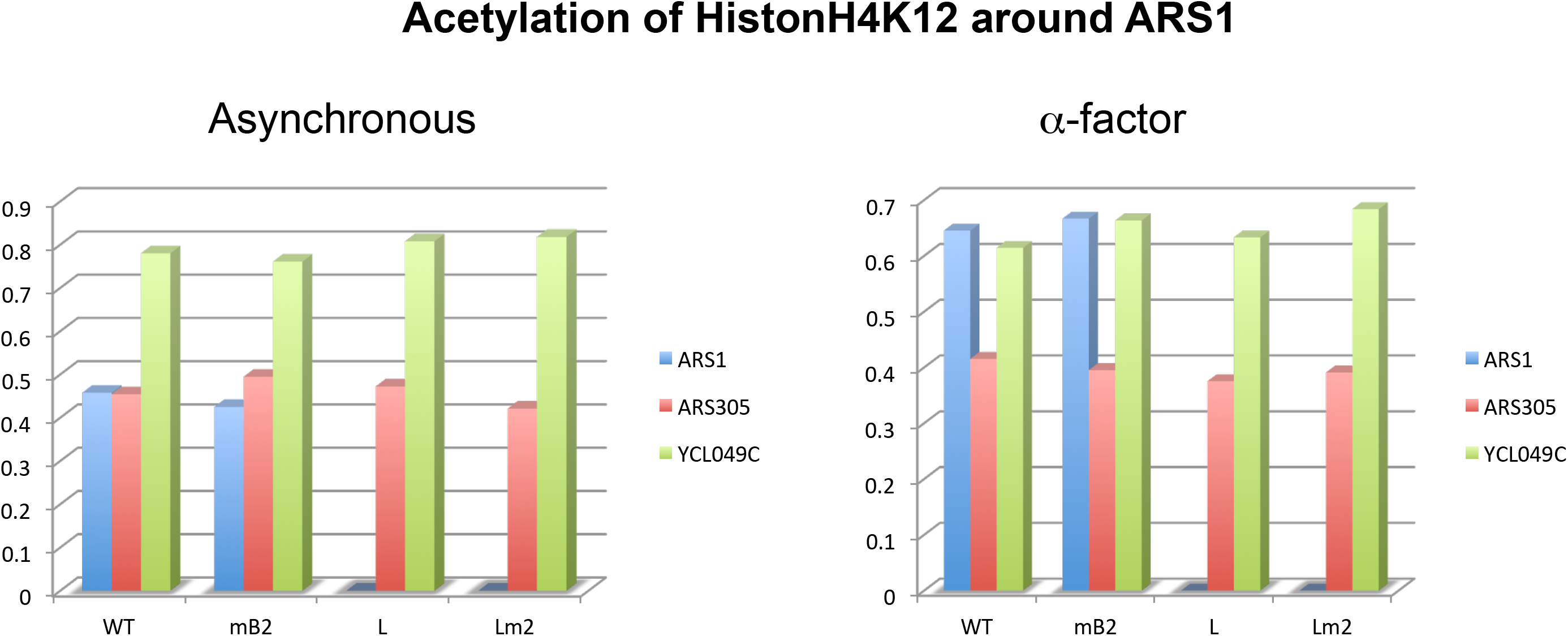
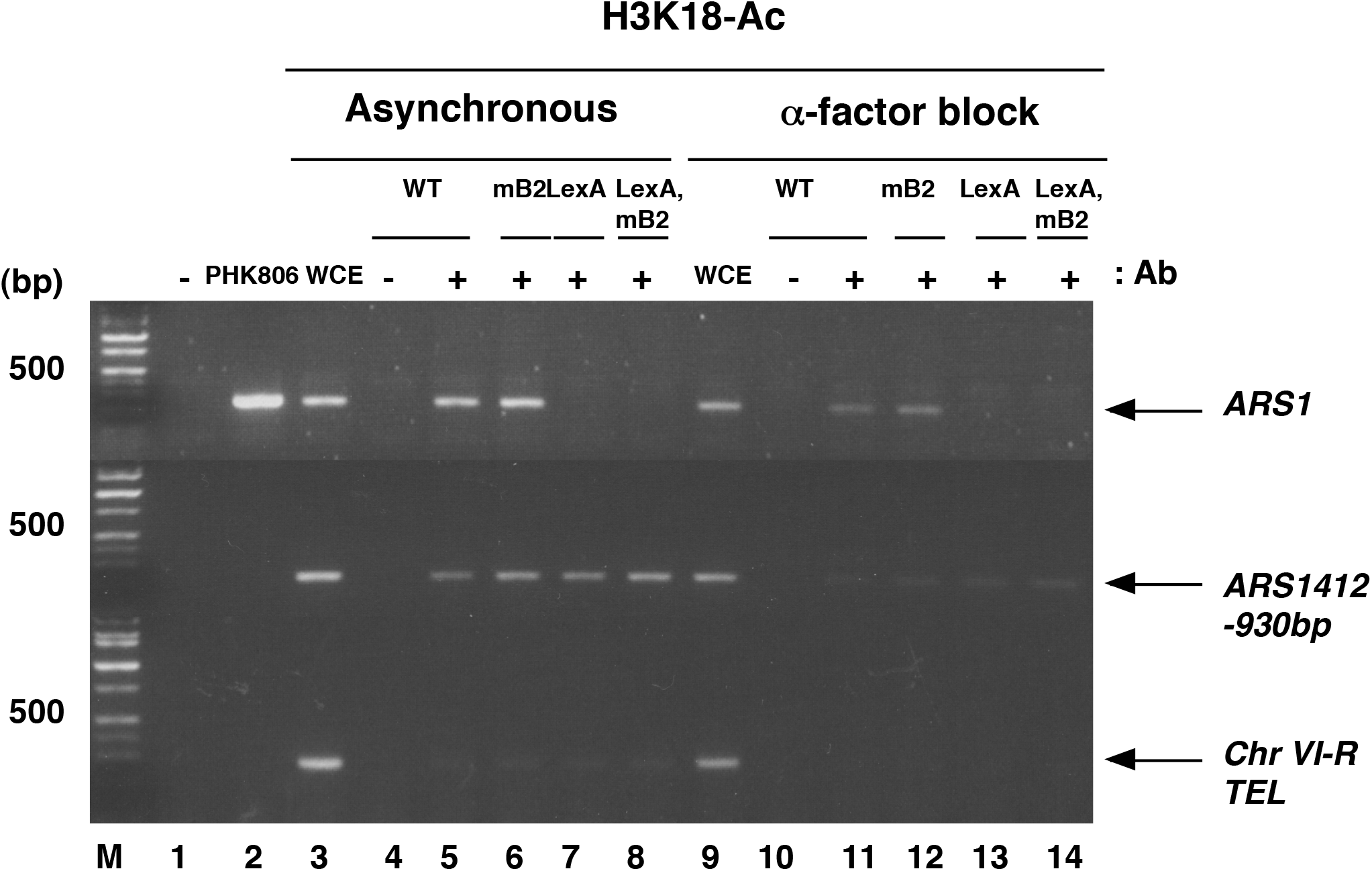
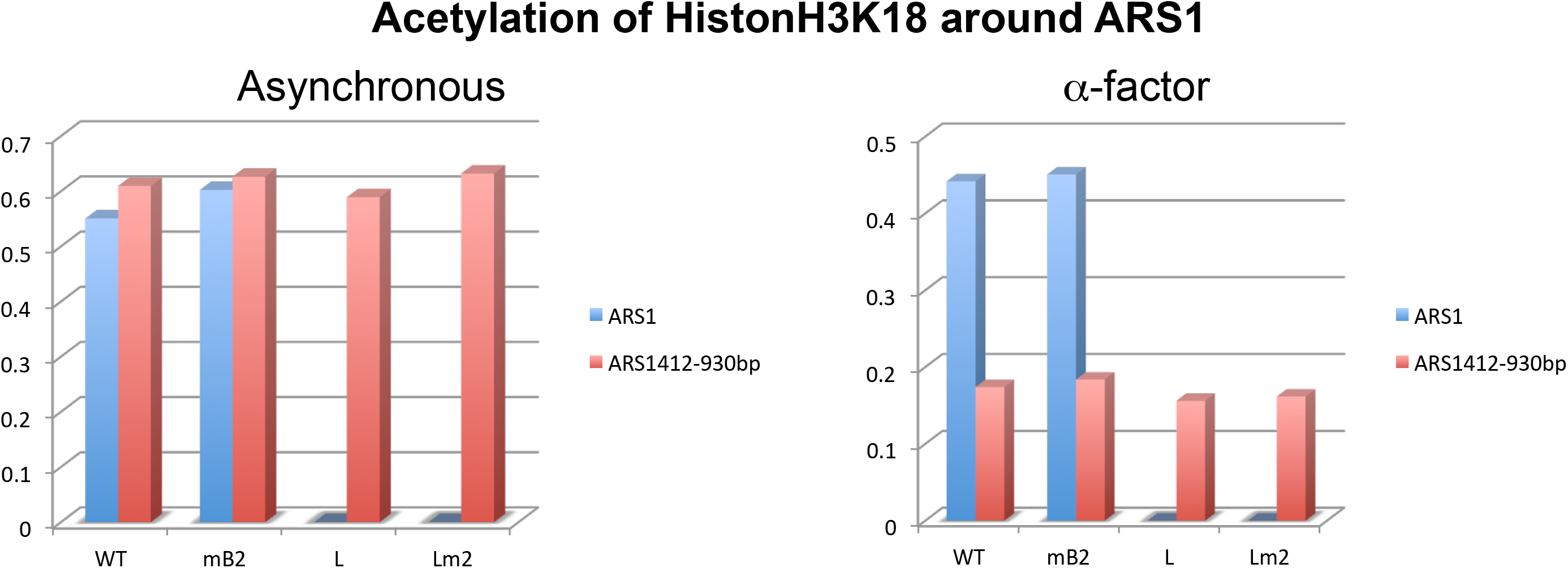
Analysis of chromatin modification surrounding wild-type and mutant *ARS1* sequences using an anti-acetylated histone H4K12 antibody and an anti-acetylated histone H3K18 antibody. (*A*) Cells harboring wild-type *ARS1* and the *ARS1* mutations indicated in the figure were either harvested as an asynchronous population, or were synchronized in late G1 with α-factor. ChIP assays were performed with the anti-acetylated histone H4K12 antibody as described in the Methods section. PCR with primers (Vogelauer et al. 2002) specific to *ARS1*, *ARS305*, and the non-origin sequences, *YCL049C* (acetylated histone H4K12) and *ChrVI-R TEL* (unacetylated histone H4K12), were used to amplify the precipitated DNA for analysis by agarose gel electrophoresis and EtBr staining. The experiments were repeated twice, with similar results. Yeast strains used were YHM08, YHK007, YHK008 and YHK009 (Table S2). M, molecular weight marker. (*B*) The amounts of ARS1 WT PCR products obtained with ChIP assay relative to that obtained from mB2, B3/LexA and B3/LexA, mB2(Lm2) but not in ARS305 of (A). (*C*) Cells harboring wild-type *ARS1* and the mutations indicated in the figure were either harvested as an asynchronized population or were synchronized in late G1 with α-factor. ChIP assays were performed with anti-acetylated histone H3K18 antibody, as described in the Methods section. PCR with primers (Vogelauer et al. 2002) specific to *ARS1, ARS305*, the non-origin sequences, *ARS1412*-930bp (acetylated histone H3K18) and *ChrVI-R TEL* (unacetylated histone H3K18), were used to amplify the precipitated DNA for analysis by agarose gel electrophoresis and EtBr staining. The experiments were repeated twice, with similar results. Yeast strains used were YHM08, YHK007, YHK008 and YHK009 (Table. S2). (*D*) The amounts of ARS1 WT PCR products obtained with ChIP assay relative to that obtained from L but not in ARS305 of (C).

Next, we focused on Esa1, a member of the MYST family of histone acetyltransferases, and Gcn5, as they are responsible for the acetylation of H4K12 and H3K18, respectively (Suka et al. 2001). Using a yeast strain expressing HA-tagged Esa1 at the same level as the endogenous Esa1 (Reid et al. 2000) (Fig. S7), we analyzed whether Esa1 and *ARS1* interact, using the ChIP assay (Fig. 3A). Esa1 bound to *ARS1* with a similar strength as it did to its other known target loci, *RPS11B*, *RPS19B, RPS8A, RPL2B*, RPL5 and *RPS5* (Reid et al. 2000) (Fig. 3A) but not to the *CYC1* and ACT1 locus.

**Figure 3.**
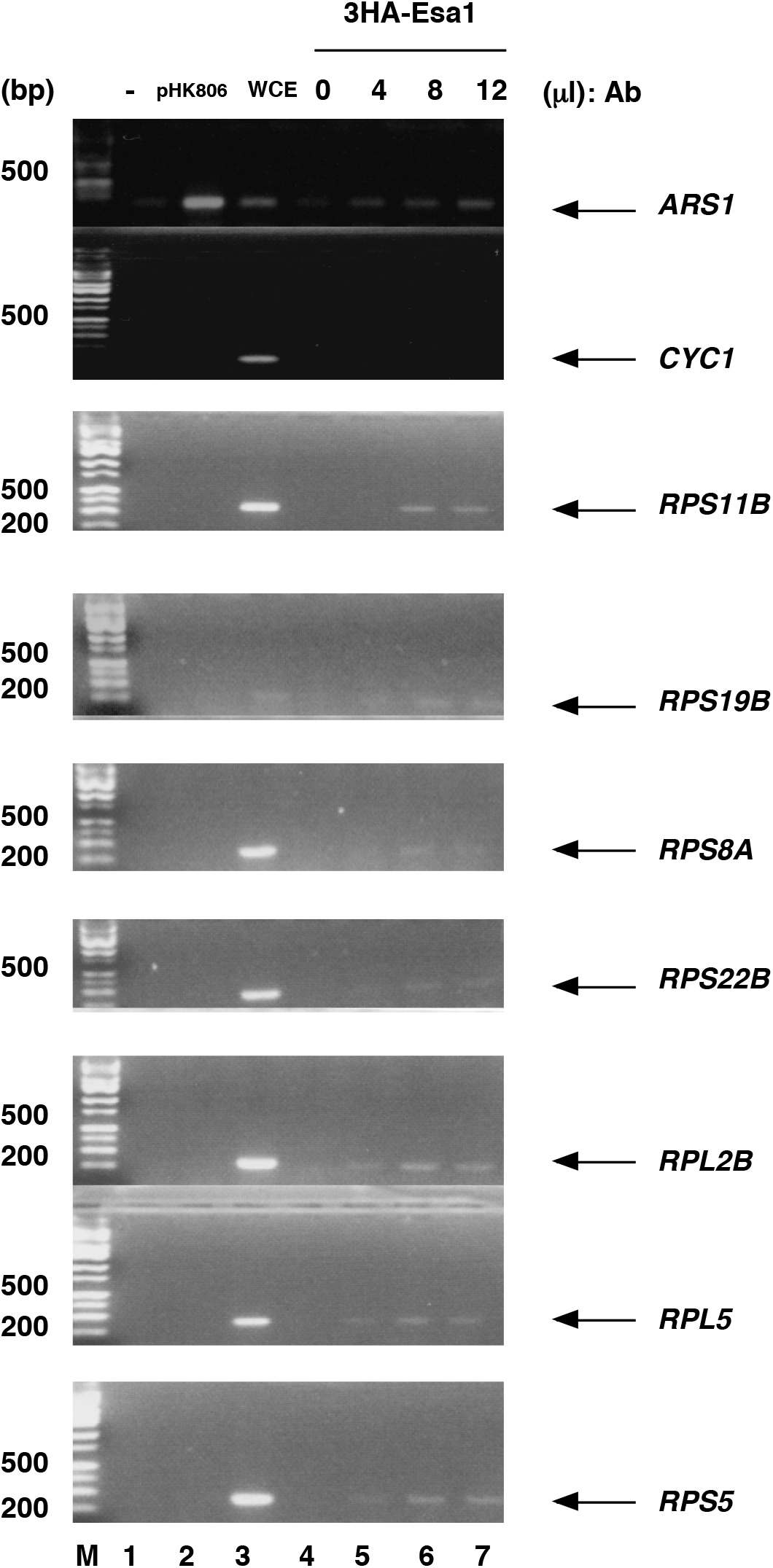
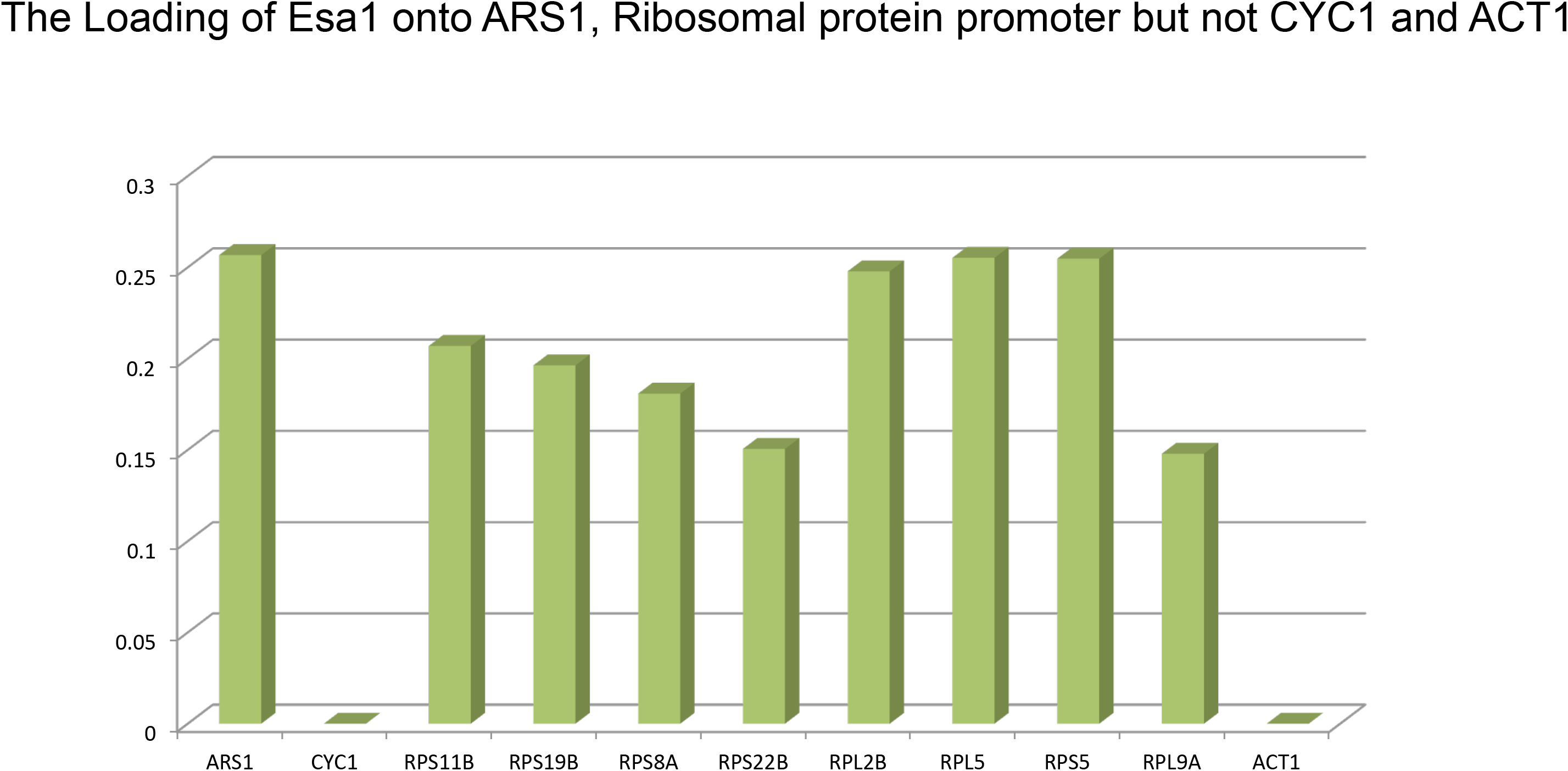
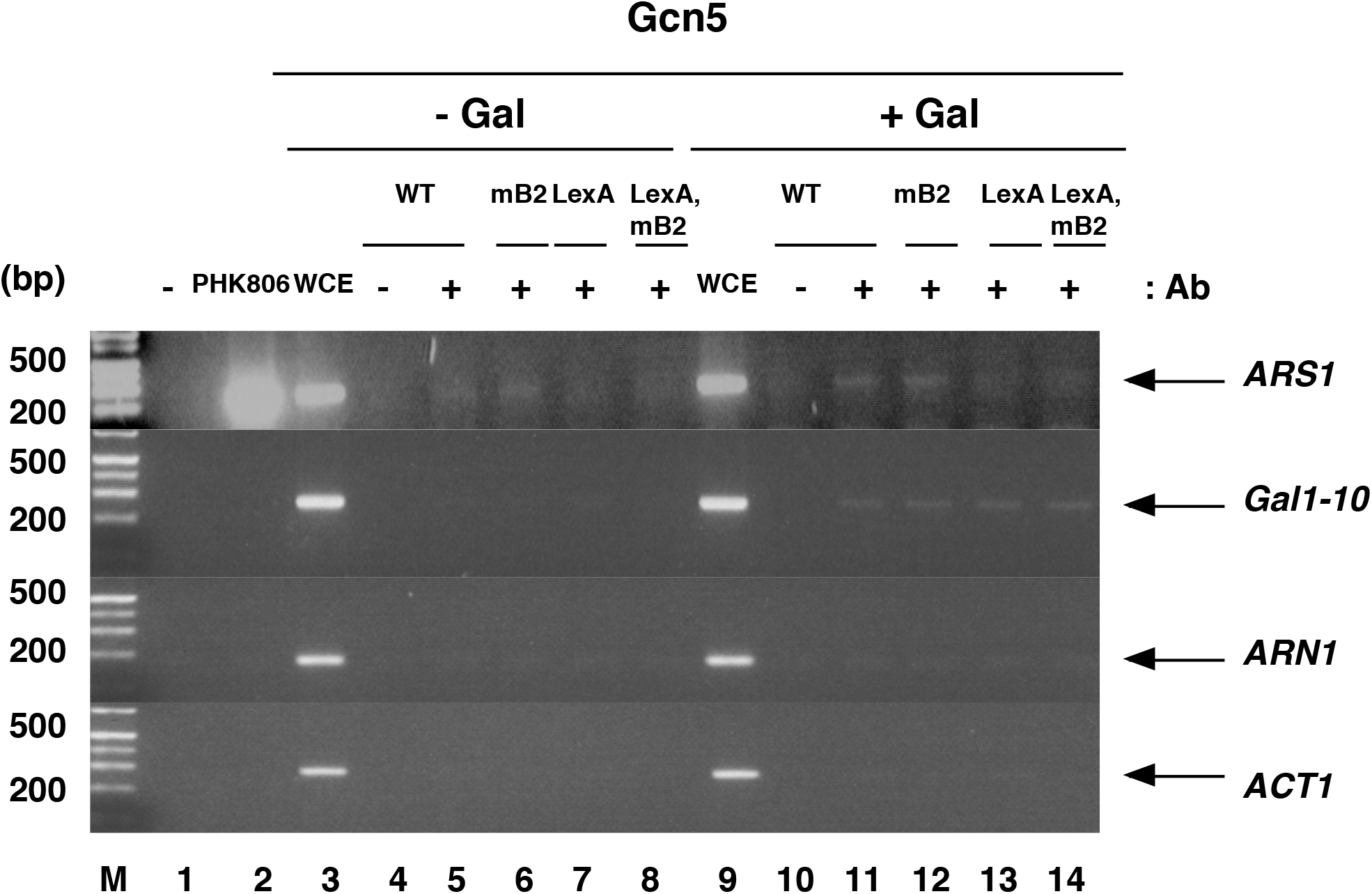
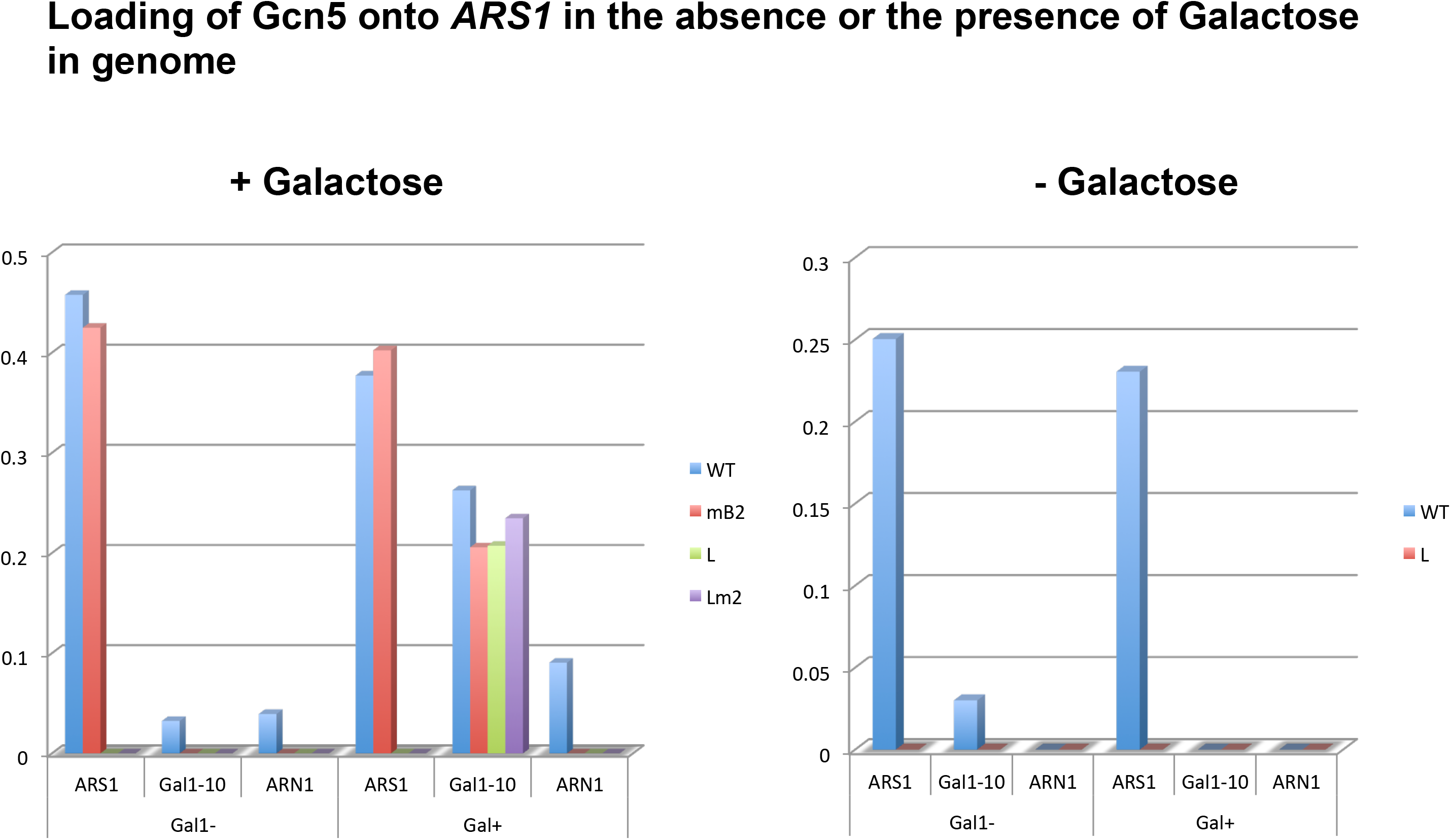
Esa1 and Gcn5 are loaded onto ARS1. (*A*) ChIP assay of 3HA-Esa1. ChIP assays were performed with anti-HA, as described in the Methods section. The indicated promoters of ribosomal proteins, ARS1 and CYC1 were detected. The experiments were repeated twice, with similar results. The yeast strain used was YHK013 (Table S2). M, molecular weight marker. (*B*) The amounts of the indicated promoters of ribosomal proteins, ARS1, CYC1 and ACT1 PCR products obtained with ChIP assay relative to that obtained from WCE. (*C*) Cells harboring wild-type *ARS1* and the mutations indicated in the figure were cultured in YPD (-Gal) or YPGal (+Gal). ChIP assays were performed with anti-Gcn5, as described in the Methods section. PCR with primers (Lemieux and Gandreu 2004) specific to *ARS1*, the non-origin sequences *Gal1-10* (to which Gcn5 is known to localize), *ACT1* and *ARN1* (to which Gcn5 never localizes) were used to amplify the precipitated DNA for analysis by agarose gel electrophoresis and EtBr staining. The experiments were repeated twice, with similar results. Yeast strains used were YHM08, YHK007, YHK008 and YHK009 (Table S2). M, molecular weight marker. (*D*) The amounts of ARS1 PCR products obtained with ChIP assay relative to that obtained from WCE in the presence and the absence of Galactose in C.

Similarly, the ChIP assay, using an anti-Gcn5 antibody, indicated that Gcn5 also localized to *ARS1*, irrespective of galactose induction, which is known to cause Gcn5 to bind to its target loci, such as *Gal1-10* (Fig. 3B; Fig. S8) (Lemieux and Gandreu 2004). Gcn5 was not detected at other loci, including *ARN1* or *ACT1* (Fig. 3B, lanes 5 and 11; Fig. S8, lanes 5 and 9). The B3 mutation reduced the binding of GCN5 with *ARS1* to undetectable levels (Fig. 3B, lanes 7 and 13; Fig. S8 lanes 6 and 10), but it did not affect the localization at *Gal1-10*. These data showed that Gcn5 constitutively bound to *ARS1* in a B3-dependent manner.

Since HBO1, a mammalian counterpart of Esa1, was shown to interact with ORC in mammals (Iizuka and Stillman 1999; Burke et al. 2001), we investigated the interaction between ORC and Esa1 using an immunoprecipitation assay. When Orc2-9myc was immunoprecipitated with anti-myc (Fig. S7B lanes 5 and 6) or an anti-ORC antibody (Fig. S7B lanes 8 and 9), HA-Esa1 was co-precipitated, indicating that Esa1 binds to ORC. These data suggest that a complex consisting of Esa1 and ORC may be responsible for acetylation of H4K12 at the ARS elements.

Immuno-precipitation assay using an anti-myc antibody and strains expressing Gcn5-9myc indicated that Gcn5-9myc was co-precipitated with endogenous Abf1 (Fig. S7B, lanes 2-4 compared with lanes 10-12). Conversely, when Abf1 was precipitated by an Abf1-specific antibody, Gcn5-9myc could be detected in the precipitates (Fig. S7C, lanes 6-8 compared with lanes 14-16). These data suggest that Gcn5 is recruited to *ARS1* by the binding of Abf1 to the B3 element and Gcn5 subsequently induces the acetylation of H3K18 in the surrounding chromatin.

We also analyzed genetic interaction between B2 or B3 element and g*cn5, esa1 or* histone deacetylase *rpd3*. Rpd3 is a histone deacethylase, which deacetylates all four core histones in both a global and targeted manner (Shahbazian and Grunstain, 2007). As shown in Table.S4, the B3 element showed synthetic defects *gcn5* or rpd3 but not *esa1*. On the other hands, the B2 mutant showed synthetic defect with *esa1* but not *gcn5 or rpd3*. These data suggests that the level of histone acetylation regulates the ARS activity and GCN5, RPD3function through B3 element, while ESA1 functions through B3 and B2 element. Our data suggested that the chromatin around origins was modified dynamically during DNA replication, which is also suggested by Grunstein et al.

To test the function of Gcn5 in DNA replication, we analyzed the stability of a plasmid bearing the Gcn4-binding site instead of the B3 element (pARS1 (B3/GCN4). The transcriptional activator Gcn4 is known to target Gcn5 to specific sites (Kuo et al. 2000). As shown in Fig.S15, pARS1 (B3/GCN4) was more stable than the B3 mutant (B3/LexA), though less stable than the wild-type *ARS1* plasmid. This suggests that Gcn4 can partially replace Abf1 function, probably by recruiting Gcn5. We were unable to perform the stability assay using *Δgcn5* cells, because the *Δgcn5* strains could not maintain the wild-type *ARS1* plasmid (Table S2). This also suggests that *gcn5* is important for ARS plasmid maintenance. In addition, the replacement of B3/LexA to B3/GCN4 recovered the maitotic stability defects in several strains especially in K6447 (using T testing, p<0.05)(Fig.S13). Consistent with the stability assay described above, ChIP analysis indicated that Orc2 localized to pARS1 (B3/Gcn4) at the same level as the wild-type *ARS1*, but did not localize to pARS1 (B3/LexA or B3/mB3) (using T testing, p<0.05) (Fig. 4A; Fig. S14). We found that H3K18 was acetylated around ARS1 (B3/GCN4) (Fig. S6; Fig. 4B, lane 6), but H4K12 was not (Fig. 4C, lane 6). Thus, Gcn4/Gcn5 appears to recruit ORC to the ARS, but at levels that are too low to recruit Esa1 for H3K18 acetylation.

**Figure 4.**
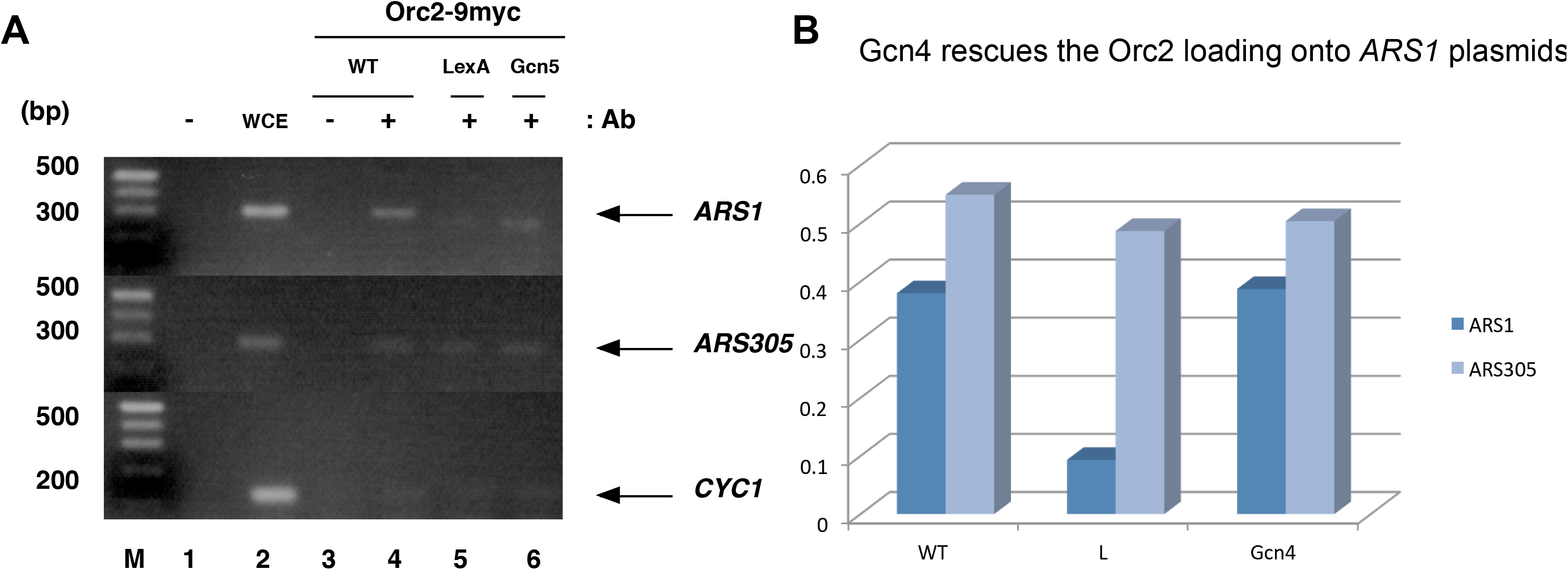
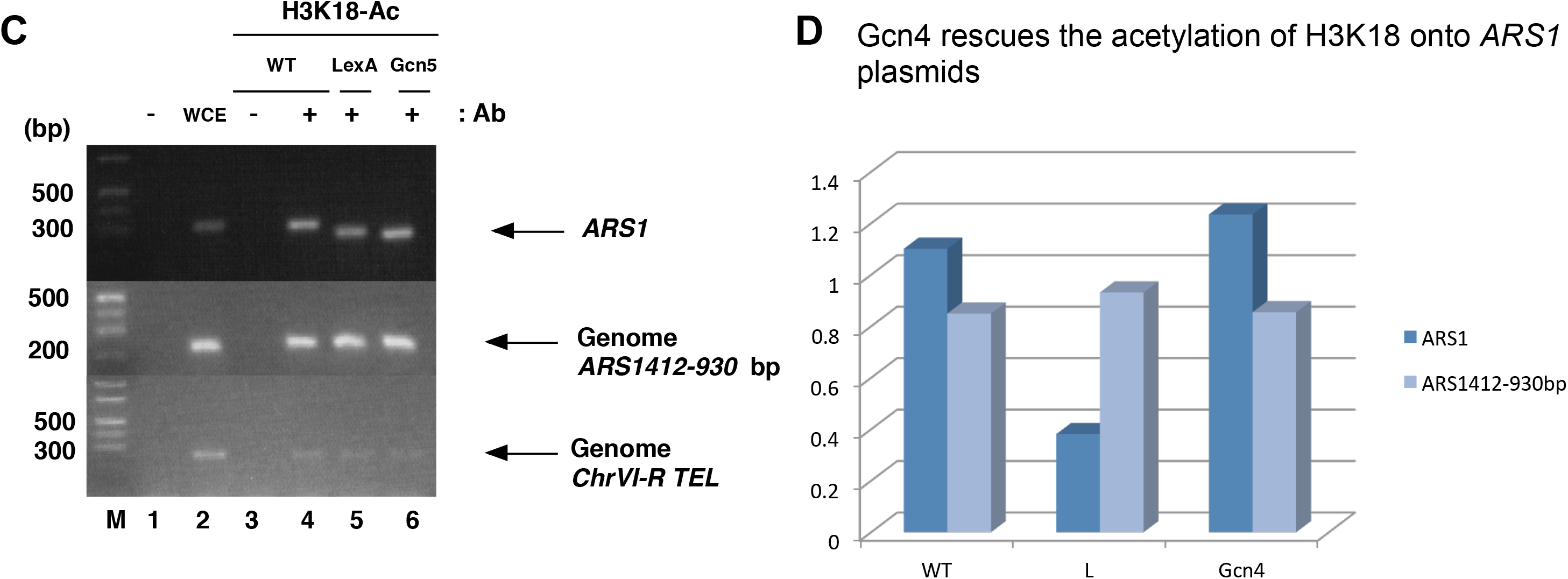
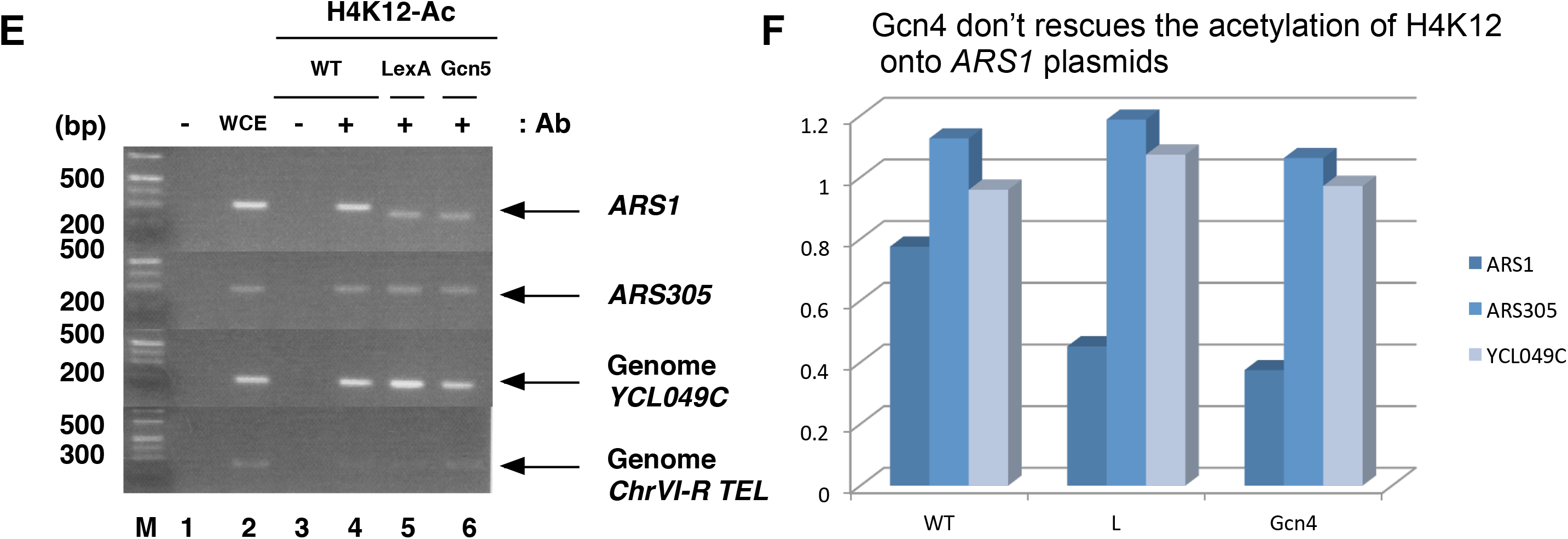
Partial rescue of the B3 mutation in *ARS1* by Gcn4. (*A*) Plasmid ChIP assay of the Gcn4 site. The yeast strain K6447, harboring the plasmids indicated in the figure, was grown in SCM-Ura+Ade for the mitotic stability assay. ChIP assays were performed with anti-myc, as described in the Methods section. PCR with primers specific to the pARS1 plasmid, *ARS305* and *CYC1*were used to amplify the precipitated DNA for analysis by agarose gel electrophoresis and EtBr staining. M, molecular weight marker. (*B*) The amounts of ARS1 PCR products obtained with ChIP assay relative to that obtained from L and Gcn4 but not in ARS305 in A. (*C*) Plasmid ChIP assay of the Gcn4 site. The yeast strain K6447, harboring the plasmids indicated in the figure, was grown in SCM-Ura+Ade for the mitotic stability assay. ChIP assays were performed with anti-acetylated histone H3K18, as described in the Methods section. PCR with primers specific to the pARS1 plasmid, *ARS1412*-920bp and *ChrVI-R TEL* were used to amplify the precipitated DNA for analysis by agarose gel electrophoresis and EtBr staining. M, molecular weight marker. (*D*) The amounts of ARS1 PCR products obtained with ChIP assay relative to that obtained from L and Gcn4 but not in ARS305 in C. (*E*) Plasmid ChIP assay of the Gcn4 site. The yeast strain K6447, harboring the plasmids indicated in the figure, was grown in SCM-Ura+Ade for the mitotic stability assay. ChIP assays were performed with anti-acetylated histone H4K12, as described in the Methods section. PCR with primers specific to the pARS1 plasmid, *ARS305*, *YCL049* and *ChrVI-R TEL* were used to amplify the precipitated DNA for analysis by agarose gel electrophoresis and EtBr staining. The experiments were each repeated twice, with similar results. (*F*) The amounts of ARS1 PCR products obtained with ChIP assay relative to that obtained from L and Gcn4 but not in ARS305 in E.

## Discussion

In this study, we showed that a transcription factor ABF1 binding to B3 element in ARS1 is required for efficient Orc loading onto ARS1. Abf1 recruits a HAT, GCN5, resulting in histone acetylation including H3K18 around ARS1. Therefore, we speculate that the change of chromatin structure induced by histone acetylation by GCN5 promotes loading of ORC (Fig S9). This speculation is further supported by the specific synthetic defects between *orc* mutations (*orc2-1*, *orc1-3* and *orc1-4*) and B3 element mutations (mB3, B3/Gal4 and B3/LexA)

Lipford and Bell suggested that ORC could still bind to *ARS1* in the B3 mutant, based on their findings with an in vivo nucleosome mapping technique that is rather indirect assay to see the ORC binding (Lipford and Bell 2001), which showed that a B3 mutation had no effect on the nucleosome positioning pattern around the A element, which represents the binding site of the ORC. However, Hu et al. showed that a mutation in the A element itself did not change the overall nucleosome positioning (Hu et al. 1999), indicating that ORC binding status does not affect the nucleosome positioning. Since it has been shown that a B3 mutation results in decreased, but still significant, replication activity of *ARS1*, both on the plasmids and the chromosomes (Marahrens and Stillman 1992; Marahrens and Stillman 1994; Kohzaki et al. 1999), functional ORC binding to ARS1 seems to be maintained, to a certain extent, in the B3 mutant. We speculate that the absence of the B3 element/Abf1 binding destabilizes the ORC/A element association and that this can be detected by the ChIP assay, but not by the nucleosome mapping assay. In addition, the nature of the mutation introduced into the B3 element might also be responsible for the different results. In a previous report, it was suggested that Orc1 does bind to a B3-deficient *ARS1*(Aparicio et al.1997). However, in that study, an XhoI linker was substituted for the B3 element in the ChIP assay, while in this present study, we substituted the LexA binding site for the B3 element. In support of our assumption, we found that the extent of the synthetic growth defects in *orc* mutants bearing B3 mutations was influenced by the type of B3 mutation present; Lex A substitution mutants showed more severe defects compared with other mutations (B3/Gal4 and B3/mB3) (Fig. S4). In addition, the difference in the ORC subunit used in the ChIP assay, or the conditions of cross-linking, may have affected the sensitivity of the ChIP assay. The decreased association of other replication components (Mcm, Rfa, Cdc45, DNA polymerase 2(Pol2) and DNA polymeraseα-primase) with the B3-deficient ARS1 chromosome and plasmid (Fig. 1 and Fig. S3) further supports the requirement of the B3 element for ORC loading.

In this study, we have shown that Abf1 recruits another HAT, Gcn5, to ARS1 in order to acetylate histones (including H3K18) around *ARS1*. Therefore, we speculate that the change of chromatin structure induced by Gcn5-directed histone acetylation promotes the loading of ORC. Several lines of evidence have indicated, directly or indirectly, that chromatin structure regulates DNA replication; for example, histones located around the origins that control amplification of the chorion gene loci in *Drosophila* follicle cells are hyperacetylated (Aggarwal and Calvi 2004). Tethering the histone deacetylase Rpd3 to the amplification origin decreased its replication activity, whereas tethering the Hat1 homologue, Chameau acetyltransferase, increased origin activity (Aggarwal and Calvi 2004; Kohzaki and Murakami in preparation). Therefore, we assume that Gcn5, which is recruited by Abf1 to the B3 element, alters chromatin structure by acetylating histones around *ARS1*, thereby stabilizing or enhancing the association with ORC.

We also found that Esa1 was recruited to *ARS1* by ORC. Esa1, a yeast histone acetyltransferase of the MYST family and its mammalian counterpart HBO1, has been linked to the ORC. HBO1 associates with pre-RC components, such as Orc1, Orc2, Mcm2, Geminin and Cdc6, *in vitro* and regulates the initiation of DNA replication by acetylating the pre-RC component (Iizuka and Stillman 1999; Burke et al. 2001; Iizuka et al. 2006). Sas2 has been genetically linked to the Orc2 and Orc5 subunits of ORC, though it is not yet clear whether it plays a role in regulating replication initiation (Ehrenhofer et al.1997). Recently, Kurat et al showed Gcn5 and Esa1 each contribute separately to maximum DNA synthesis rates(Kurat et al, 2017).

We speculate that Esa1, recruited by ORC, acetylates histones and probably also the components of the pre-RC to facilitate subsequent initiation steps.

The use of specific origins appears to change dramatically during development (Kohzaki and Murakami 2005; Aladjem 2007). At the onset of zygotic transcription in *Drosophila* and *Xenopus*, replication origins become restricted to a limited set of sites. Also, eukaryotic replication origins exhibit different initiation efficiencies and activation timings. It is possible that the alteration of chromatin structure, induced by transcription factors, could be one of the mechanisms that regulate the timing of replication and origin selection. It is noteworthy that, in mammals, many transcription factors are also proto-oncogenes, such as c-Jun, E2F, Rb, c-Myb and c-Myc. Though, the oncogenicity of these genes is thought to be due to their involvement in the disruption of transcription, it is possible that their malfunction in the regulation of DNA replication would undoubtedly contribute to their oncogenic potential (Kohzaki and Murakami 2005; Dominguez-Sola et al. 2007). More detailed information about the inter-relationship between transcriptional regulation and DNA replication would need to be stored to understand the whole process of cell growth regulation and oncogenesis.

## Methods

The Supplemental Methods section provides detailed information regarding all experimental procedures: (1) yeast strains and culture; (2) mitotic stability assay; (3) synchronization; (4) ChIP assay; (5) plasmid and yeast construction; (6) protein expression and (7) imunoprecipitation and western blotting.

## Acknowledgements

We dedicate this work to Dr. Yasuo Kawasaki (Osaka University, Osaka, Japan) and Dr. Yamaguchi-Iwai Yuko (Kyoto University, Kyoto, Japan). We do not know what we would have done without them during the early stages of this work. We thank B. Stillman, H. Araki, Y. Kawasaki, H. Maki, L. Graudreau, K. Struhl and J.F.X. Diffely for the yeast strains and the plasmids used in this study. We are also grateful to A. Abiko, S. Nishino, K. Kamei, S. Hara, T. Uemura and M. Sugita for helpful discussions. This work was partially supported by the Japanese Leukemia Research Fund. H.K. was supported by a KIT VL grant and Y.M. was supported by a Grant-in-Aid for Scientific Research on Priority Areas from the Japan Society for the Promotion of Science.

